# FennoTraits: Dataset of plant functional traits and community composition in northern European flora

**DOI:** 10.64898/2026.04.07.716889

**Authors:** Pekka Niittynen, Julia Kemppinen

## Abstract

We present here FennoTraits, which is a dataset of plant functional trait and community composition data which we collected from Fennoscandia across northern Finland, Norway, and Sweden in 2016-2025. This dataset has 42 049 abundance estimations and 155 794 functional trait observations from 10 traits representing 373 vascular plant species collected from 1 235 study sites within seven study areas. The trait measurements consist of size-structural, leaf economic, leaf spectral, and reproductive traits. The species represent the majority of the native vascular plant species that occur at the seven study areas, and many of the species occur in all seven areas across the two biomes and their ecotone: tundra and boreal forests. Each study area has distinct characteristics and a range of habitats: tundra, meadows, wetlands, shrublands, and boreal forests. These areas are under low anthropogenic influence, and many of the sites are within protected areas that are reserved for nature conservation and scientific research. Finally, we provide with this dataset a general description of the main trait patterns and profiles of the northern European flora.

## Introduction

Plant functional traits can reveal the mechanisms driving ecosystem dynamics and species interactions. Functional traits are thus a powerful tool and have emerged into a critical research topic across ecology, biogeography, and environmental sciences (Violle et al. 2007, Diaz et al. 2016, Jennifer L. Funk et al. 2017). Plant functional traits are measurable plant characteristics that influence the growth, survival, and reproduction of plants, and also plant interactions with the environment and other organisms (Pérez-Harguindeguy et al. 2013). Therefore, it is essential to understand how and why functional traits vary across and within species (Kemppinen and Niittynen 2022, Laughlin 2024). Variation in functional traits can explain how plants adapt to changing environmental conditions, such as climate change, and how plants contribute to ecosystem services, such as carbon sequestration (Gerlinde B. De Deyn et al. 2008, Christiane Roscher et al. 2012, Georges Kunstler et al. 2016). Ultimately, functional diversity is a key component of biodiversity, contributing to ecosystem resilience and stability (Cadotte et al. 2011, Mammola et al. 2021, Carmona et al. 2021).

Northern European ecosystems are facing rapid warming due to anthropogenic climate change, which is challenging the resilience of these ecosystems and their provided services (Cohen et al. 2014, Rantanen et al. 2022). Northern ecosystems play significant roles in carbon storage regulation and nature-based livelihoods, and they also harbor unique biodiversity (Hobbie et al. 2000, Ford et al. 2021). Northern Europe encompasses Finland, Sweden, and Norway, representing ecosystem diversity ranging from northern boreal forests to sub-Arctic and oro-Arctic tundra. These ecosystems are characterised by strong environmental gradients due to their rich geodiversity, providing mosaics of habitats and vegetation types that are adapted to cold and wet climates (Wielgolaski 1975, Austrheim and Eriksson 2001, Kuuluvainen and Aakala 2011). The northern boreal forests are primarily composed of coniferous species Norway spruce (*Picea abies*) and Scots pine (*Pinus sylvestris*). Towards higher latitudes, the coniferous forest shifts to sub-Arctic mountain birch forest (*Betula pubescens* ssp. *czerepanovii*). Towards higher altitudes, the deciduous forest transitions to dwarf-shrub dominated oro-Arctic heath (chiefly, *Empetrum nigrum* and *Betula nana*), and finally, to mostly barren mountain tops.

Measuring plant functional traits across wide environmental gradients is important for investigating how plants are shaped by environmental change. Ultimately, functional trait data can be leveraged in establishing more effective conservation strategies and restoration efforts, which requires extensive trait measurements (Carlucci et al. 2020). Some plant functional traits are informative but laborious, and in turn, expensive to measure, such as root traits. Whereas, other traits are more cost-efficient to collect, enabling replication of a high number of plant species, communities, and study sites across large gradients. Such cost-efficient traits include plant height, leaf area, specific leaf area (SLA), and leaf dry matter content (LDMC). These are relatively fast and easy to measure, and require only a ruler, scale, scanner, and oven (Figure 1) (Pérez-Harguindeguy et al. 2013). Plant height and leaf area represent size-structural traits, and SLA and LDMC represent leaf economic traits, and together these four traits form the two principal trait variation axes globally, and are thus most often used in ecological research (Diaz et al. 2016).

**Figure 1.**
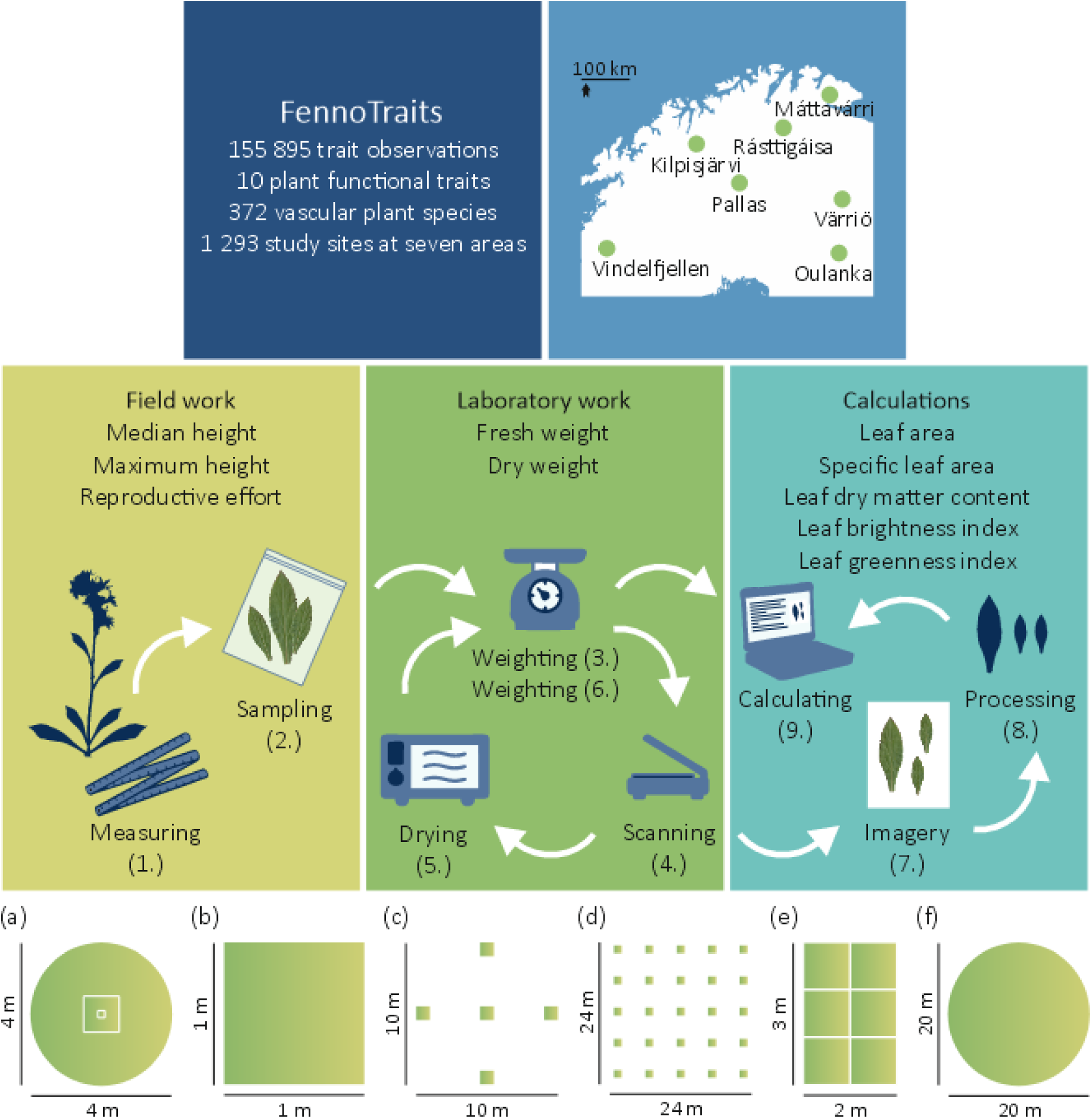
Sampling protocol and study designs (a-f). We present a comprehensive dataset of plant functional trait measurements that we collected from Fennoscandia across northern Finland, Norway, and Sweden in 2016-2025.

Here, we present FennoTraits, which is a taxonomically and spatially comprehensive dataset of plant functional trait measurements in Fennoscandia across Finland, Norway, and Sweden (Figure 1). We collected this dataset primarily from protected areas at seven study areas in 2016-2025. With this documentation of northern European flora, we aim to advance the understanding of biodiversity, trait variability, and ecological responses across different environmental conditions and large gradients. The dataset includes 42 049 abundance estimations from 373 vascular plant species and 155 794 functional trait observations from 10 traits representing 1 235 study sites, allowing analyses on intraspecific trait variability and its responses to a range of environmental gradients. The dataset consists of plant community composition with a nested plot structure, enabling analyses on community structure, diversity, and mechanisms linked to the locally measured plant traits. Lastly, this dataset is unique in the sense that it was collected and processed only by two researchers, maximising data comparability across the many study designs and study areas. We followed the best practices for open and reproducible science in planning, collecting, documenting, and publishing this dataset (Pérez-Harguindeguy et al. 2013, Hampton et al. 2015, Wilkinson et al. 2016, Jenkins et al. 2023).

## Methods

### Study areas

We collected the data at seven study areas (Figure 1; Table 1; see Supporting information for maps Figures S1-S7), four in Finland, two in Norway, and one in Sweden. We chose these study areas due to their floristic diversity, accessibility and on-going research collaboration. Many of the study areas are nearby research stations, facilitating both the field and laboratory work.

**Table 1.**
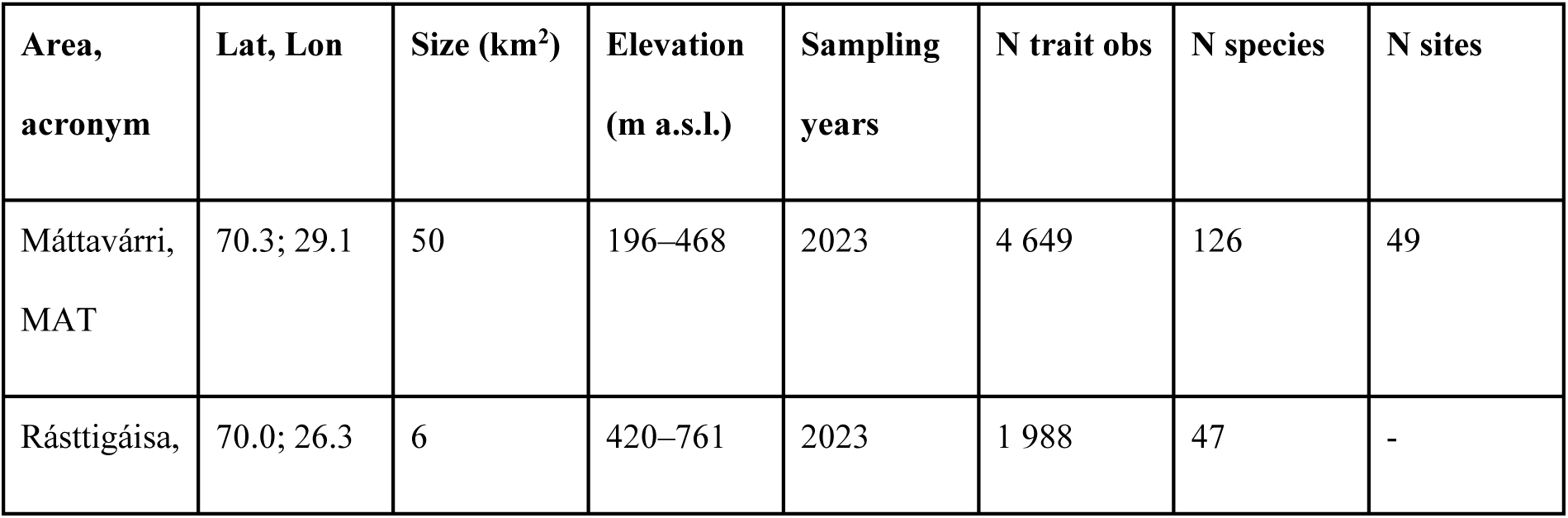
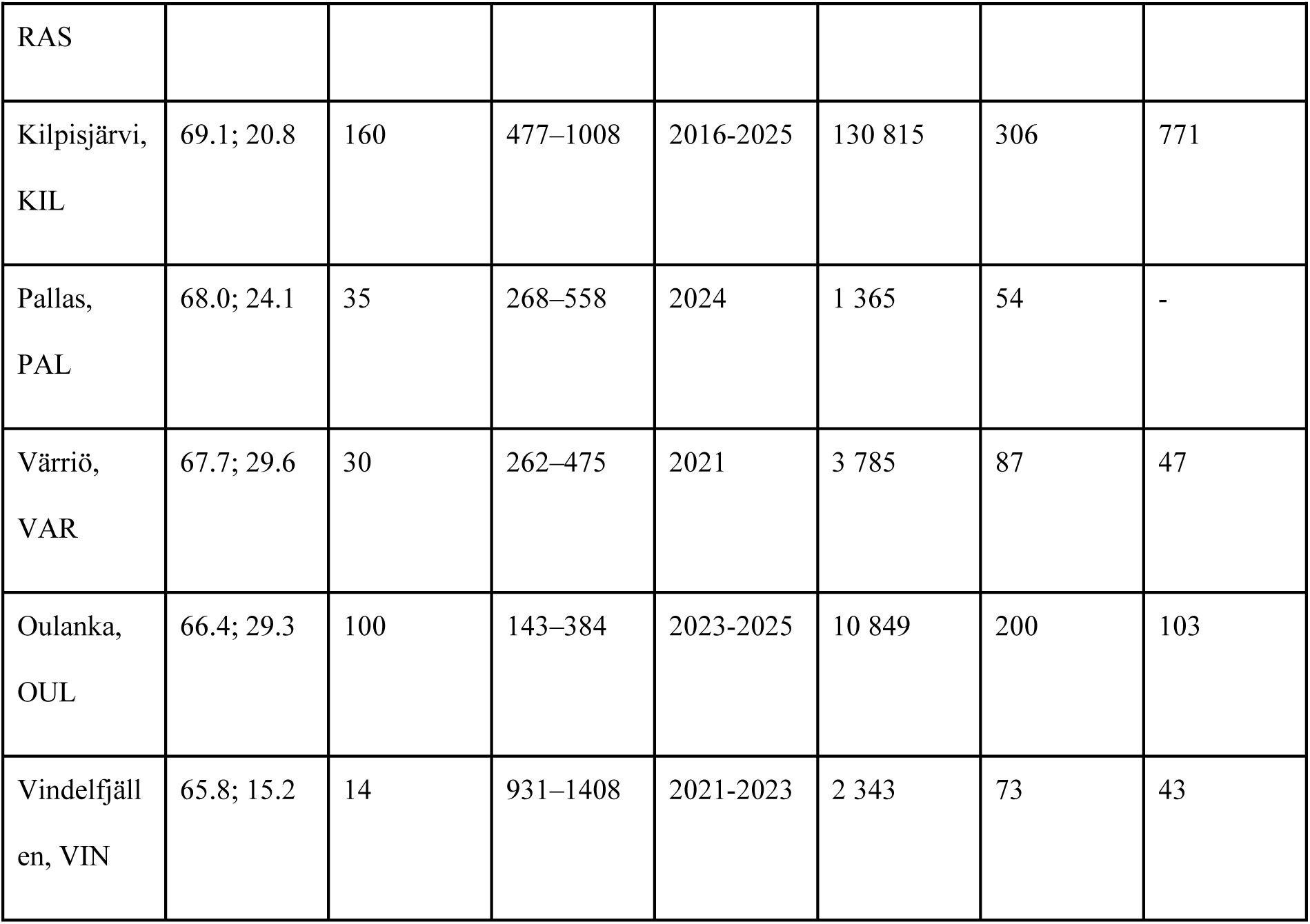
Summary statistics of the study areas. N trait obs = number of unique trait observations; N species = number of unique species in data; N sites = number of unique study sites with full plant community surveyed.

### Máttavárri area

The Máttavárri area is located in northern Norway. In this low-Arctic area, the dominating vegetation type is dwarf shrub tundra. The summits are bare.

### Rásttigáisá area

The Rásttigáisá area is located in northern Norway, close to the Norway-Finland border. In this sub-Arctic area, the dominating vegetation type is dwarf shrub tundra. The summits are bare.

### Kilpisjärvi area

The Kilpisjärvi area is located in north-western Finland, partly extending to the Salloaivi mountain close to the border of Finland and Norway. In this sub-Arctic area, the dominating vegetation type is dwarf shrub tundra and mountain birch forest. The summits are bare. The area is diverse and heterogeneous with both acidic and calcareous bedrocks. The area has several conservation areas, including Malla Strict Nature Reserve and Saana Nature Reserve.

### Pallas area

The Pallas area is located in north-western Finland. In this hilly northern boreal area, the dominating vegetation type is coniferous forest, which forms the tree line. The summits are tundra heath vegetation. Aapa mires are common in the lowlands. The area is within the Pallas-Yllästunturi National Park.

### Värriö area

The Värriö area is located in north-eastern Finland, on the Finland-Russia border. In this northern boreal area, the dominating vegetation type is coniferous forest, which nearly reaches the mountain tops. The summits are tundra heath vegetation. Aapa mires are common in the lowlands. The area is within the Värriö Strict Nature Reserve.

### Oulanka area

The Oulanka area is located in eastern Finland, on the Finland-Russia border. In this boreal area, the dominating vegetation type is mixed forest, which covers the entire area except the open wetlands and small patches of herb-rich deciduous forests. The area is within the Oulanka National Park.

### Vindelfjällen area

The Vindelfjällen area is located in northern Sweden, close to the Sweden-Norway border. In this mountain area, the dominating vegetation type is mountain tundra heath and extensive herb-rich snowfields. The summits are bare. The area is within the Vindelfjäll Nature Reserve.

## Study designs

At the seven study areas, we used different study designs, and in the Kilpisjärvi area we had 11 different study designs (Figure 1a-f; Table 2). Across all study areas and designs, the plant functional trait and plant community composition data are fully comparable because we used the same methods and observers. Yet, each study design was established for a specific purpose, and therefore, the spatial, environmental, and temporal structures differ among the areas and designs. Where possible, we have provided references to the original study designs for detailed information on the design and related environmental data. We have provided coordinates to the entire dataset with GPS accuracy, and at most sites, we used a high-accuracy Global Navigation Satellite System (chiefly, GeoExplorer GeoXH 6000 Series; Trimble Inc., Sunnyvale, CA, USA; or a comparable device) that provides up to centimeter-scale positioning accuracy. For many of the study designs, we used random stratification to select the study sites. We used the random stratification to pre-select a set of candidate sites, maximising the coverage of the main environmental gradients within each study area. The variables that we used for stratifying the environmental space vary among the study areas and designs. We provide references for each design when possible, and in short, the variables included, e.g., total canopy cover, deciduous canopy cover, distance to forest edge, altitude, potential incoming solar radiation, and topographic wetness.

**Table 2.** The data from the Kilpisjärvi study area originated from 11 different study designs. N trait obs = number of unique trait observations; N species = number of unique species in data; N sites = number of unique study sites with trait or abundance data.

| Study design | Acronym | Sampling years | N trait obs. | N species | N sites |
| --- | --- | --- | --- | --- | --- |
| Woody plant design | KIL_WOO | 2016-2017 | 9679 | 21 | 223 |
| Saana-Jehkas gradient design | KIL_MI | 2017-2025 | 16084 | 141 | 228 |
| Malla gradient design | KIL_MAL | 2020-2025 | 6966 | 136 | 70 |
| Rare Arctic design | KIL_RA | 2020-2025 | 23862 | 247 | 182 |
| Ailakkavaara gradient design | KIL_AIL | 2021-2025 | 4874 | 129 | 41 |
| Arthropod sampling design | KIL_ROS | 2021 | 4665 | 114 | 35 |
| Microclimate grid design | KIL_ITV | 2021 | 4107 | 6 | 148 |
| Spring design | KIL_L | 2022-2023 | 4169 | 128 | 32 |
| Geodiversity gradient design | KIL_X | 2023-2025 | 4451 | 141 | 39 |
| Community experiment design | KIL_EXP | 2023-2025 | 48184 | 139 | 144 |
| Seasonal monitoring design | KIL_SEA | 2024 | 1551 | 11 | 3 |

At many of the study designs, we used a nested plot structure (Figure 1a), unless we mention otherwise (Figure 1b-f). The nested plot structure means that at each site we had three plot scales nested so that the centre of the plots align. The plot scales and forms are: 0.2 m x 0.2 m squared plot, 1.0 m x 1.0 m squared plot, and 2.0 m radius circular plot (Figure 1a).

In addition to the study designs (Figure 1a-f), we also collected extra leaf samples opportunistically in the field (coded as “EXT” in the leaf trait data). We targeted these efforts to gain more trait data for species that were underrepresented in the functional trait data collected at the studied plots. This means that the plant functional trait data also contains leaf trait measurements from these extra leaf samples, without accompanying height trait measurements or the plant community composition data.

The study sites and plots are primarily unmanipulated, except for the leaf trait sampling, unless we mention otherwise, for instance, the Kilpisjärvi: Community experiment design.

### Máttavárri: Gradient design

In the Máttavárri area, we sampled plant functional trait and plant community composition data in 2023. This design consists of 49 study sites, and here, we used the nested plot structure (Figure 1a).

### Rásttigáisá: Gradient design

In the Rásttigáisá area, we sampled plant functional trait data in 2023. This design consists of 49 study sites with 1 m x 1 m study plots (Figure 1b). These data consist only of leaf traits of the most abundant plant species in the plant communities. This means that the data do not include height measurements or the plant community composition data. The original study design is described in detail in Rissanen et al. (2023).

### Kilpisjärvi: Woody plant design

In the Kilpisjärvi area, we used a design focusing on woody plant species in tundra. In this design, we measured woody species cover and height in 2016-2017. This design consists of 223 study sites with five study plots, each 1 m x 1 m. In total, we had 1 053 plots which we placed hierarchically, so that at each site, one plot was at the centre of the site and four plots were placed in the four cardinal compass directions five meters from the centre plot (Figure 1c). These data consist of species-specific height measurements (median, maximum) and cover percentages of woody plant species. This means that the data do not include other trait measurements or the plant community composition data. The original study design is described in detail in Kemppinen et al. (2021b) and Kemppinen et al. (2018).

### Kilpisjärvi: Saana-Jehkas gradient design

In the Kilpisjärvi area, we sampled plant functional trait and plant community composition data in 2017-2025. This design consists of 228 study sites with 1 m x 1 m study plots. We have studied 50 of these sites more intensively, and used here the nested plot structure (Figure 1a) and annual resampling of leaf traits. This design overlaps with the centre plots of the Kilpisjärvi Woody plant design. The original study design is described in detail in Kemppinen et al. (2021b) and Tyystjärvi et al. (2022).

### Kilpisjärvi: Malla gradient design

In the Kilpisjärvi area, we sampled plant functional traits and plant community composition in 2020-2025. This design consists of 70 study sites, and here, we used the nested plot structure (Figure 1a). The original study design is described in detail in Aalto et al. (2022) and Kemppinen et al. (2023).

### Kilpisjärvi: Rare Arctic design

In the Kilpisjärvi area, we used a design focusing on rare Arctic vascular plant species and thus targeted their habitats, such as calcareous heaths and meadows. In this design, we sampled plant functional trait and plant community composition data in 2020-2025. This design consists of 182 study sites, and here, we used the nested plot structure (Figure 1a).

### Kilpisjärvi: Ailakkavaara gradient design

In the Kilpisjärvi area, we sampled plant functional trait and plant community composition data in 2021-2025. This design consists of 41 study sites, and here, we used the nested plot structure (Figure 1a). The original study design is described in detail in Aalto et al. (2022) and Kemppinen et al. (2023).

### Kilpisjärvi: Arthropod sampling design

In the Kilpisjärvi area, we used a design focusing on arthropods. In this design, we sampled plant functional trait and plant community composition data in 2021. This gradient design consists of 35 study sites, and here, we used the nested plot structure (Figure 1a). The sites were used for sampling arthropods using malaise traps and ground traps, however, these were not placed in the plots. The original study design is described in detail in Peña-Aguilera et al. (2023).

### Kilpisjärvi: Microclimate grid design

In the Kilpisjärvi area, we used a design focusing on within-species microclimate relationships of six common tundra vascular plant species. In this design, we sampled plant functional trait data in 2021. This design consists of six study grids with 25 study plots, each 1 m x 1 m. In total, we had 150 plots which we placed in a grid layout, so that in each grid, the 25 plots were placed at six m intervals (Figure 1d). These data consist only of plant functional traits of the six species. This means that the data do not include the plant community composition data. In two plots, the focal species were not present and thus we measured traits from 148 plots. The original study design is described in detail in Kemppinen & Niittynen (2022).

### Kilpisjärvi: Spring design

In the Kilpisjärvi area, we used a design focusing on springs. In this design, we sampled plant functional trait and plant community composition data in 2022-2023. This design consists of 32 study sites at or around springs, and here, we used the nested plot structure (Figure 1a).

### Kilpisjärvi: Geodiversity gradient design

In the Kilpisjärvi area, we used a design focusing on geodiversity. In this design, we sampled plant functional trait and plant community composition data in 2023-2025. This design consists of 39 study sites at locations with pronounced geomorphological processes (e.g., cryoturbation and fluvial activity), and here, we used the nested plot structure (Figure 1a).

### Kilpisjärvi: Community experiment design

In the Kilpisjärvi area, we used a design with a plant community experiment. In this design, we sampled plant functional trait and plant community composition data in 2023, retrieving a baseline data prior to the community manipulations that followed in 2024 and 2025 when trait sampling and community surveys were also repeated. This design consists of 24 replicated study grids (each 2 m x 3 m) with six 1 m x 1 m study plots (Figure 1e). In total, we had 144 plots that represent herb-rich meadow vegetation. Within a grid, we had one control plot and the rest had a different manipulation: 1) species with the highest LDMC removed; 2) species with the lowest LDMC removed; 3) tallest species removed; 4) shortest species removed; 5) species removed randomly. We conducted the manipulations in a given plot community by active removals (i.e., cutting the above-ground parts) so that at least half of the total vascular plant cover was removed. The removals were repeated 2-3 times per growing-season.

### Kilpisjärvi: Seasonal monitoring design

In the Kilpisjärvi area, we used a design focusing on seasonality of intra-specific trait variation of 11 common tundra vascular plant species. In this design, we sampled leaf functional trait data in 2024. This design consists of three study sites with 10 m-radius circular study plots (Figure 1f). We conducted a weekly sampling for 15 weeks, the entire growing season. These data consist only of the leaf functional traits of the 11 species. This means that the data do not include height measurements or the plant community composition data. The original study design is described in detail in Niittynen et al. (2026).

### Pallas: Gradient design

In the Pallas area, we sampled plant functional trait data in 2024. This design consists of 23 study sites with 1 m x 1 m study plots (Figure 1b). These data consist only of leaf traits of the most abundant plant species in the plant communities. This means that the data do not include height measurements or the plant community composition data. The larger original study design is described in detail in Lehtinen et al. (2025).

### Värriö: Gradient design

In the Värriö area, we sampled plant functional trait and plant community composition data in 2021. This design consists of 47 study sites, and here, we used the nested plot structure (Figure 1a). The original study design is described in detail in Aalto et al. (2022) and Kemppinen et al. (2023).

### Oulanka: Gradient design

In the Oulanka area, we sampled plant functional trait and plant community composition data in 2023-2025. This design consists of 103 study sites, and here, we used the nested plot structure (Figure 1a).

### Vindelfjällen: Gradient design

In the Vindelfjällen area, we sampled plant functional trait and plant community composition data in 2021-2023. This design consists of 43 study sites with 1 m x 1 m study plots (Figure 1b).

## Taxonomy

We identified the plants to species level always when it was possible. There were only few exceptions: 1) The genus *Taraxacum* is known for its complex taxonomy, and therefore, we identified it only to genus level; 2) *Alchemilla* species can be difficult to identify to species level in field, and therefore, in some of our subdatasets it is only at genus level. An exception to *Alchemilla* spp. is *A. alpina*, which we always identified at species level, because it is easy to identify due to its separated leaflets. We used the Leipzig Plant Catalogue as the backbone for our taxon nomenclature. We harmonised the taxon names using the *lcvplants* R package (Freiberg et al. 2020). Hybrids are common in *Salix* and *Carex* genera. When we suspected a hybrid, we named the taxon after the most likely pair of species that formed the hybrid. In the dataset, all hybrids are easy to separate using the provided taxonomic rank of each recorded taxon.

## Plant community composition data

We collected the plant community composition data from the plots by identifying all vascular plant species and visually estimating their species-specific cover percentages. The sum of cover values within plots may exceed 100 %, as plants often overlap each other.

In the community data, there are four species (total of 10 occurrences in the plant community composition data) that are classified as sensitive species in Finland. Therefore, we anonymised these species in the entire dataset (i.e., SpeciesA-D) to protect their precise locations in Finland, following the guidelines of the Finnish Biodiversity Information Facility (FinBIF).

## Plant functional trait data

We collected the plant functional trait data by following the protocol outlined in Kemppinen & Niittynen (2022) and Niittynen et al. (2026) which are based on the handbook for standardised measurements of plant functional traits (Cornelissen et al. 2003, Pérez-Harguindeguy et al. 2013). We collected data on 10 plant functional traits (Table 3), namely, median height, maximum height, reproductive effort, fresh weight, dry weight, leaf area, SLA, LDMC, leaf brightness index (BITM; Equation 1), and leaf greenness index (Excess Green index; ExG; Equation 2). The height traits are measured at plot-level, which means that a species can have multiple height measures per site in those study designs in which we used the nested plot structure. We measured the rest of the traits at species- and site-level, which means that we measured or sampled several individuals from each focal species and pooled the samples by site. Below, we explain our 9-step plant functional trait protocol in chronological order (Figure 1.1-9).

**Table 3.** Summary of the 10 plant functional traits and their prevalence in the data.

| Trait | Unit | Explanation | N trait obs. | N species | N sites |
| --- | --- | --- | --- | --- | --- |
| median height | cm | Estimated median vegetative height (inflorescence excluded) of a focal species within a plot. | 27201 | 327 | 1157 |
| maximum height | cm | Maximum vegetative height (inflorescence excluded) of a focal species within a plot. | 27200 | 327 | 1157 |
| reproductive effort | unitless | Intensity of the sexual reproduction effort of a species at a study site. From 0 to 5, 0 indicating no signs of sexual reproduction effort (flowers, berries, fruits), 3 indicating that half of the individuals show efforts of sexual reproduction, and 5 indicating that all individuals show efforts on sexual reproduction. | 18653 | 325 | 754 |
| fresh weight | g | Water-saturated fresh mass of the leaf | 11814 | 350 | 1039 |
| dry weight | g | Dry mass of the oven-dried leaf | 11805 | 350 | 1039 |
| leaf area | cm<sup>2</sup> | Area of the leaf derived from scanned leaf images taken of the fresh leaf | 11826 | 350 | 1040 |
| specific leaf area, SLA | mm<sup>2</sup>/mg<sup>-1</sup> | Specific leaf area (SLA) is the ratio of dry weight and leaf area. | 11809 | 350 | 1039 |
| leaf dry matter content, LDMC | g/g | Leaf dry-matter content (LDMC) is the oven-dry mass of a leaf, divided by its water-saturated fresh mass | 11814 | 350 | 1039 |
| Brightness index, BITM | unitless | An spectral index about the brightness of the upper side of the leaf (Equation 1) | 11836 | 350 | 1040 |
| Excess Green Index, ExG | unitless | A spectral index indicating the greenness of the upper side of the leaf (Equation 2) | 11836 | 350 | 1040 |

### Fieldwork

We measured plant height (Figure 1.1) with a ruler to centimeter precision (millimeter precision for the shortest plants), recording the median and maximum vegetative heights of the focal species and excluding inflorescences. The median height is a visual estimation of the typical height of the canopy of a given species within the plot. In the nested plot structure, we measured plant heights from the two smaller plots, but not from the largest circular plot size (Figure 1a).

We estimated reproductive effort (Figure 1.1) by documenting the sexual reproductive effort of the focal species. We used a scale from 0 to 5, where 0 means that we did not observe any signs of sexual reproduction efforts at the study site (i.e., flowers, berries, fresh seed capsules), and 5 means that all individuals showed signs of sexual reproduction efforts. If the individuals were large and hard to separate (e.g., *Empetrum nigrum*), 5 indicated exceptionally high density of flowers or berries. Therefore, the reproductive effort should be used as an index that indicates the relative intensity of sexual reproductive effort in a form that is comparable across species and sites. We did not estimate reproductive effort for ferns. The reproductive effort data can also be missing in cases when we were not able to determine if sexual reproduction was present. For instance, if the plants were still developing or if the trees were too tall for us to reach.

We collected leaf samples (Figure 1.2) from or near the plots using the plant community composition data to determine which species to sample at each site. In general, we sampled the entire community at a given site, except for species with a low coverage ≤ 1%. The exceptions to this rule are: 1) all protected species, because we did not sample them in the countries in which they were protected, and 2) specific study designs, where we did not collect any leaf samples (e.g., Kilpisjärvi: Woody plant design) or focused only on a set of focal species (e.g., Kilpisjärvi: Microclimate grid design). We selected fully opened and developed leaves (i.e., not curled) that showed no signs of damage, such as pathogens or herbivory. In general, we sampled one leaf from 3-4 individuals of the focal species at a given study site at each sampling time point. The exceptions to this rule were species with very small leaves, such as *Empetrum nigrum*. For those small-leaf species, we sampled 10-15 leaves per individual and three individuals per given study site at each sampling time point. Regarding all species, we pooled the sampled leaves at the species and site level to reduce the workload. Regarding evergreen shrub species, we selected leaves that were from previous years instead of new leaves that had emerged during the sampling season. We transported the leaf samples (or branches of e.g., *Empetrum nigrum*) from the field to the laboratory in zip-lock bags with a drop of water to keep the leaves fresh or to rehydrate them. In the laboratory, we stored the sample bags at 4°C and we processed the leaves within 48 hours.

### Laboratory work

We determined fresh weight (Figure 1.3) by first preparing the leaves by removing the petioles, and then, gently patting them dry from any excess water on their surfaces. Then, we weighed the leaves using a Mettler AE 100 scale (0.0001 g precision) or a comparable scale.

We scanned fresh leaves (Figure 1.4) right after weighing using a Canon CanoScan LiDE 400 scanner (600 dpi) or a comparable scanner. We imaged the adaxial leaf surface, i.e., the sun-facing side of the leaves. Some leaves were too large to fit the scanner, so we chopped them before scanning (e.g., *Matteuccia struthiopteris*).

We determined dry weight (Figure 1.5-6) by first preparing the leaves by drying them at 70°C for 48 hours using VWR VENTI-Line ovens. Then, we weighed the leaves right after drying them using a Mettler AE 100 scale (0.0001 g precision) or a comparable scale.

### Leaf segmentation

We calculated leaf area (Figure 1.7-8) using the leaf scans. We established the following procedure for leaf segmentation and shadow removal through an iterative optimisation process. The leaf segmentation method with accompanying computer code was published in Niittynen et al. (2026). This process involved applying the spectral index based rules and thresholds to a large dataset of scans across multiple species, visually inspecting the outcomes, and then fine-tuning the used indices and threshold parameters until no significant errors or artifacts were observed. The final procedure goes as follows. First, we conducted an initial leaf segmentation by applying thresholds to the blue and red channels, removing pixels in which the values of the blue channel were >180 or the red channel <30. Next, we applied a Normalized Difference Yellowness Index (NDYI, Equation 3) threshold (< 0.13) at the image borders (20 pixels closest to the margins), excluding shadows that may occasionally appear on the image margins. Then, we converted the filtered pixels into polygons, assuming that each polygon represented an individual leaf. We discarded any polygons with an area <200 pixels to eliminate dirt on the scans. We filled any small holes within the leaf polygons which are likely artifacts using the fill_holes function from the *smoothr* R package (Strimas-Mackey 2025), with a threshold of 1 000 pixels. We also excluded polygons at the image borders (<50 pixels to the margins) to remove possible shadows of the margins.

Next, we used an iterative refinement process on individual leaf polygons, further reducing shadows on the scans. Shadows often occurred around thick leaves, and the previous procedures were insufficient to exclude these shadows. So, first, we buffered each leaf polygon with 10 pixels, and then, we cropped and masked the original RGB image to this buffered area. Next, we determined the position of the leaf, which we were able to determine based on the scanner sensor geometry that systematically casts the shadows on the same side of the leaf scans (towards the upper right corner). We leveraged this to determine the position of the leaf and to estimate potential shadows in a specific direction based on that position. First, we calculated a centerline of the leaf using the *centerline* R package (Tsyplenkov 2025), involving the creation of a skeleton of the leaf polygon and tracing a path between two ‘end points’ defined by the maximum and minimum X or Y coordinates (depending on the aspect ratio of the leaf). Then, we were able to estimate the potential shadow locations relative to the orientation of the leaf. Next, we calculated a distance raster to the centerline skeleton towards the direction where shadows were possible. The shadows gradually shift from dark to light in a known direction, so we calculated a focal Pearson correlation coefficient between the distance raster and the blue channel using a moving window of 15 x 15 pixels. Then, we could exclude a pixel as shadow if the correlation coefficient was above 0.5, the Red Chromatic Coordinate index (RCC; Equation 4) exceeded 0.2, and NDYI was less than 0.15. We considered pixels with a blue channel value greater than 200 as white background and excluded them. Then, we refined the leaf margins with a focal majority filter (5×5 kernel). Finally, we used the processed RGB images from the leaf scans to quantify leaf area.

We calculated leaf brightness index (BITM, Equation 1) and leaf greenness index (ExG, Equation 2; Figure 1.7-8) using the leaf scans. We calculated colour indices that are used in satellite-based remote sensing of vegetation. These indices can be derived from red, green, and blue wavelengths (Montero et al. 2023). We conducted all routine raster and vector processing using functions from the *terra* (Hijmans 2022) and *sf* (Pebesma 2018) R packages.

The formulas for the spectral indices were as follows:

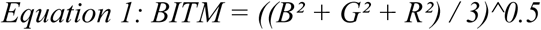

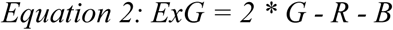

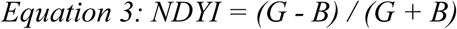

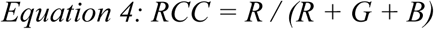

where R represents the red, G the green, and B the blue channel in the scanned RGB images.

Finally, we quantified specific leaf area (SLA; Figure 1.7-8) by calculating the ratio between leaf area and dry weight. We quantified leaf dry matter content (LDMC; Figure 1.7-8) by calculating the ratio between dry weight and fresh weight.

## Data

We provide here a general description of the main trait profiles of the northern European flora and most frequent species in the dataset (Figure 2).

**Figure 2.**
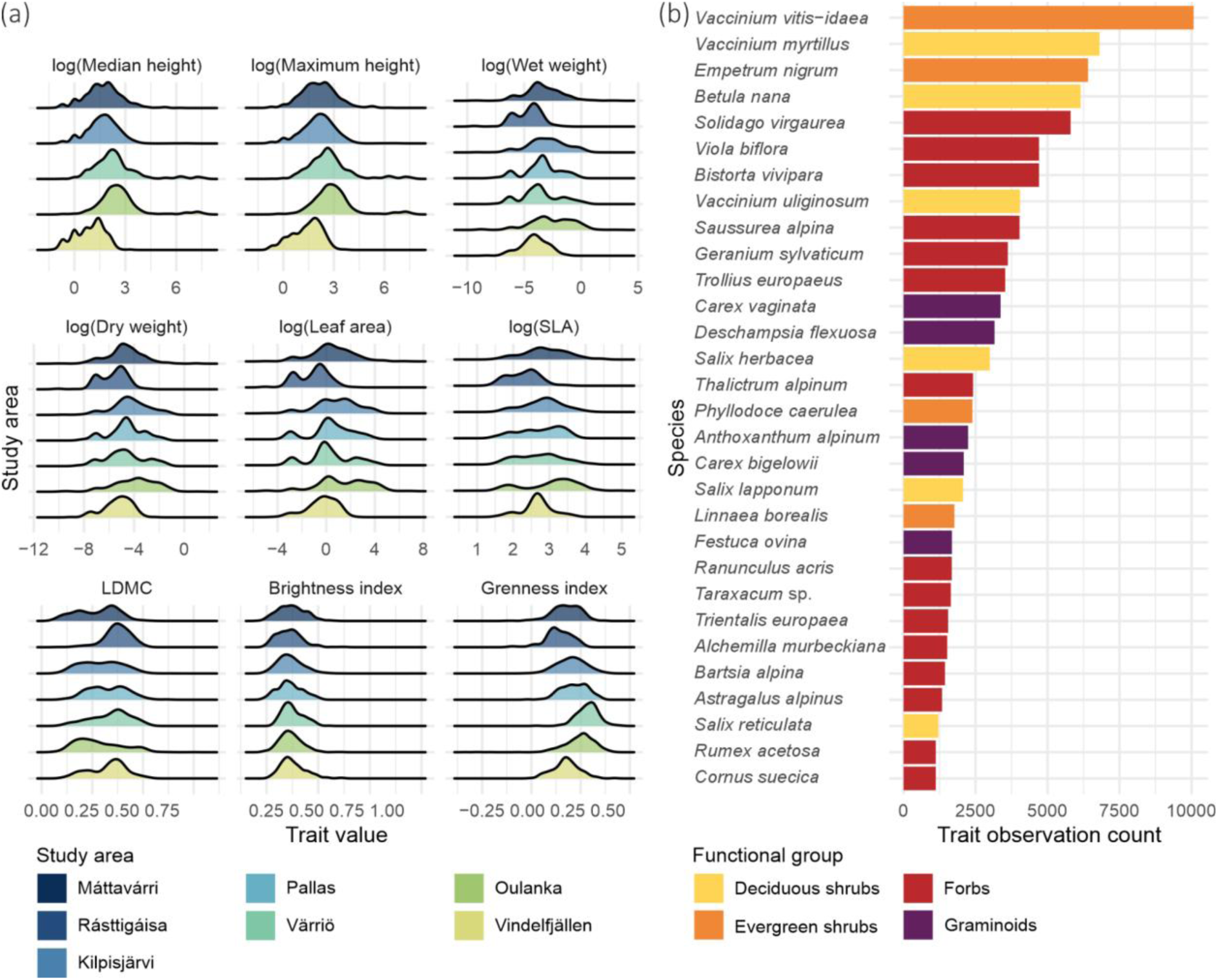
Distributions of the nine continuous traits across the seven study areas (a). The most frequent species in the plant functional trait data colored by functional group (b).

## Data and code availability

We provide the dataset in the supporting information of this article. Up-to-date version of the dataset will be maintained and openly available at a GitHub repository that we will link here after acceptance for publishing. Stable versions of the future dataset with Digital Object Identifiers (DOI) will be published in the Zenodo repository annually after major updates.

## Dataset structure and data dictionary

We provide a dataset that consists of five files: three data tables as text files, one metadata file as OpenDocument Spreadsheet, and an R script.

1. The file *FennoTraits_community_heights.csv* includes the plant community composition data and plant height data (Table 4). These data are at plot-level.
2. The file *FennoTraits_leaf_traits.csv* includes the leaf trait data and reproductive effort data (Table 5). These data are at site-level.
3. The file *FennoTraits_species.csv* is the lookup table for the observed or sampled taxa with higher taxonomy and sampling summary statistics (Table 6).
4. The file *FennoTraits_metadata.ods* includes the common metadata in three spreadsheets. This is the data dictionary of the full dataset, which includes the information in Tables 4–6.
5. The file *FennoTraits_combine_data.R* is an R script to facilitate the use of the dataset.

**Table 4.** Data dictionary for the plant community composition data and plant height data.

| Variable | Type | Description | Units / Values |
| --- | --- | --- | --- |
| site | character | Unique study site identifier, matching site in the other files | — |
| design | character | Study design the observation belongs to | — |
| area | character | Abbreviation of the study area | — |
| plot_type | character | Type of vegetation plot | a = 20cm x 20cm; b = 1m x 1m; c = circular plot with 2m radius |
| full_community | logical | Whether the record represents the full plant community (TRUE) or a focal taxon subset (FALSE) | TRUE / FALSE |
| treatment | character | Experimental treatment applied to the plot | high_LDMC = species with the highest LDMC removed; tall = the tallest species removed; low_LDMC = species with the lowest LDMC removed; random = random subset of species removed; short = the shortest species removed |
| date | date | Date of observation | YYYY-MM-DD |
| year | numeric | Year of observation | — |
| Lat | numeric | Latitude of the plot in decimal degrees (WGS84) | — |
| Lon | numeric | Longitude of the plot in decimal degrees (WGS84) | — |
| country | character | Country where the observation was made | FIN = Finland; NOR = Norway; SWE = Sweden |
| taxon | character | Taxon name as used in the dataset (may be subspecies, species, genus, or family) | — |
| cvr | numeric | Percentage cover of the taxon in the plot | % |
| median_height | numeric | Median vegetation height of the taxon in the plot | cm |
| max_height | numeric | Maximum vegetation height of the taxon in the plot | cm |

**Table 5.** Data dictionary for the reproductive effort data and leaf trait data.

| Variable | Type | Description | Units / Values |
| --- | --- | --- | --- |
| site | character | Unique study site identifier, matching site in the other files | — |
| design | character | Study design the observation belongs to | — |
| area | character | Abbreviation of the study area | — |
| treatment | character | Experimental treatment applied to the plot | high_LDMC = species with the highest LDMC removed; tall = the tallest species removed; low_LDMC = species with the lowest LDMC removed; random = random subset of species removed; short = the shortest species removed |
| date | date | Date of leaf sample collection | YYYY-MM-DD |
| year | numeric | Year of sample collection | — |
| Lat | numeric | Latitude of the observation in decimal degrees (WGS84) | — |
| Lon | numeric | Longitude of the observation in decimal degrees (WGS84) | — |
| country | character | Country where the observation was made | FIN = Finland; NOR = Norway; SWE = Sweden |
| taxon | character | Taxon name as used in the dataset (may be subspecies, species, genus, or family) | — |
| reproduction | numeric | Sexual reproductive effort index of the species at site at year of sampling | — |
| n_inds | numeric | Number of individuals sampled (and pooled) for leaf traits | — |
| n_leaf | numeric | Total (sum) of leaves sampled, pooled and processed for leaf traits | — |
| leaf_area | numeric | One-sided leaf area | cm<sup>2</sup> |
| SLA | numeric | Specific leaf area (leaf area per unit dry mass) | mm<sup>2</sup> mg<sup>-1</sup> |
| LDMC | numeric | Leaf dry matter content (dry mass per unit fresh mass) | g g<sup>-1</sup> |
| w_weight | numeric | Fresh (wet) mass of a leaf | g |
| d_weight | numeric | Dry mass of a leaf after drying | g |
| BITM | numeric | Brightness index of upper side of a plant leaves derived from scanned RGB imagery; formula = $((B^2 + G^2 + R^2) / 3)^{0.5}$ | dimensionless |
| ExG | numeric | Excess Green Index of upper side of a plant leaves from scanned RGB imagery; formula = $(2G - R - B)$ | dimensionless |

**Table 6.**
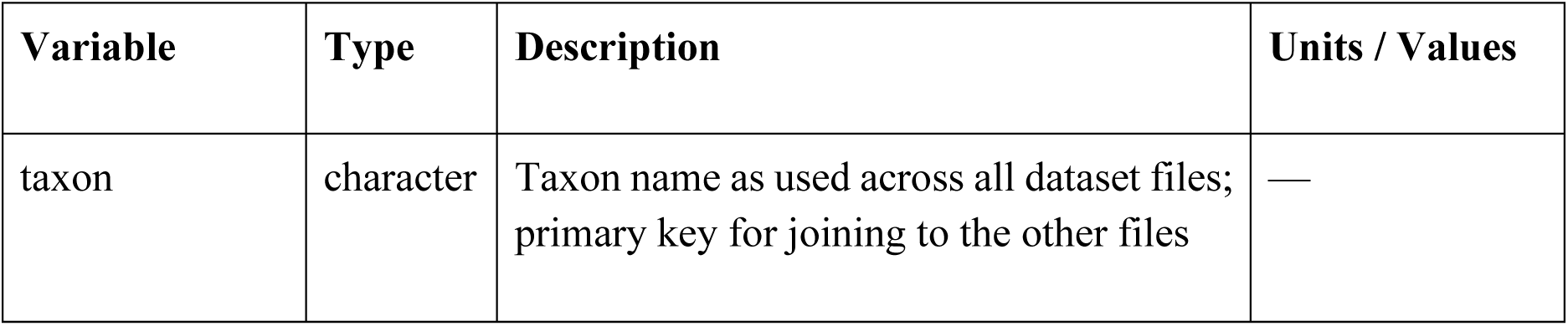
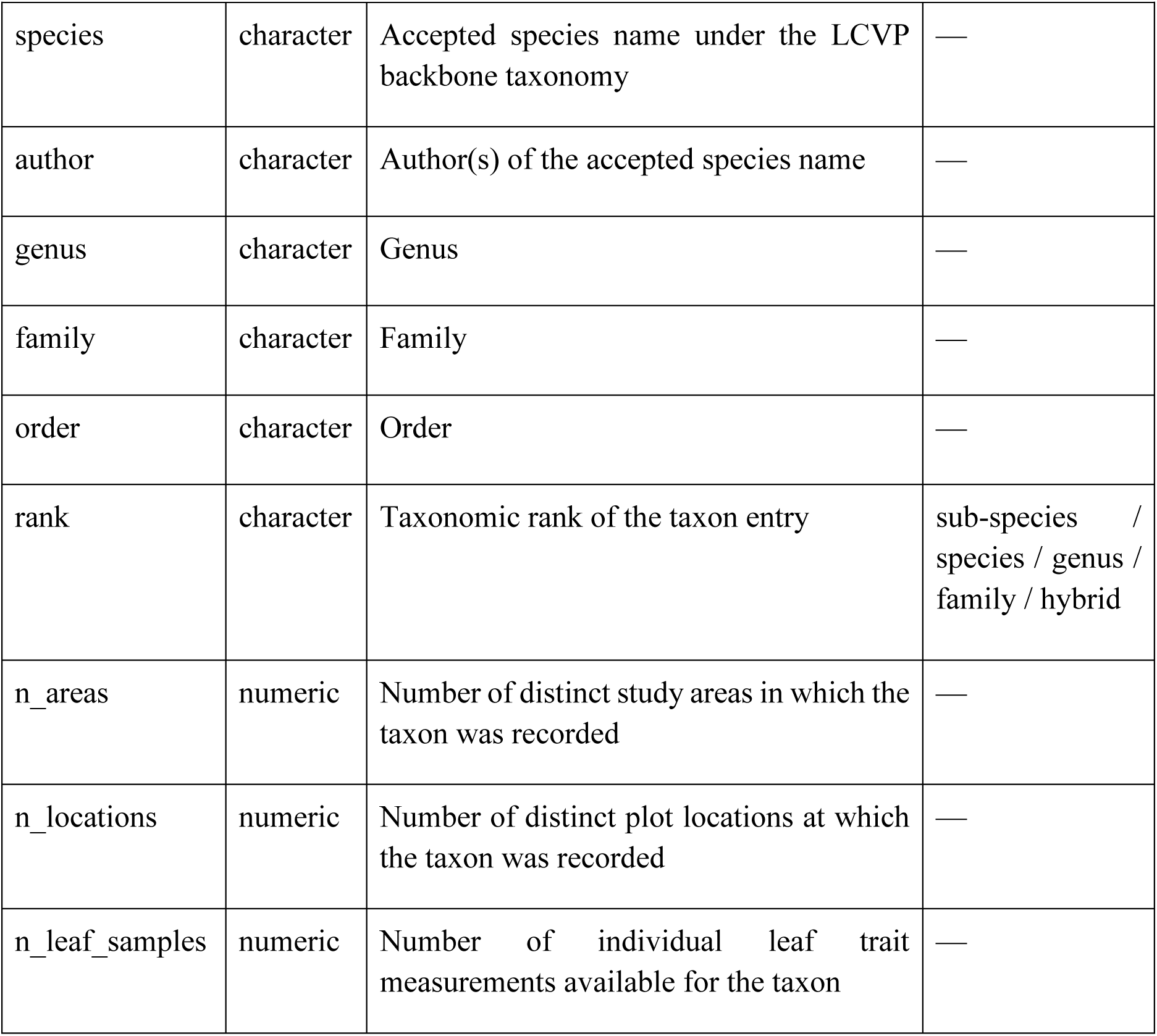
Data dictionary for the taxonomic lookup table of the observed or sampled taxa.

## Technical validation

We conducted a systematic quality control procedure on all plant community composition and plant functional trait observations. In the procedure, we combined visual inspection, biologically motivated hard thresholds, and within-species trait covariance analysis. We designed the procedure to be conservative: instead of removing observations outright based on statistical criteria alone, each step aimed to identify the specific measurement most likely to be erroneous before we set any values as missing.

First, we produced two sets of diagnostic figures for each species with sufficient data: 1) Per-species histograms of all trait distributions, allowing visual identification of extreme isolated values and measurement artefacts, and then 2) Pairwise trait-trait scatterplots with fitted linear trends for all ecologically meaningful trait pairs: log(SLA) against LDMC, log(d_weight) against log(w_weight), log(leaf_area) against log(w_weight), log(leaf_area) against log(d_weight), ExG against BITM, and log(max_height) against log(median_height). We used these plots and species-level summary statistics to identify any suspicious trait observations, which we then manually double-checked using our field notes and raw data, and finally, we corrected those observations if we found a clear source of error, such as a typing error.

After the visual inspection, we applied a set of hard bounds derived from the known biological and physical constraints of each measured variable. We flagged observations with the following criteria: 1) Cover values outside the range 0.25–100%; 2) Plant height values <0 or >3000 cm; 3) Observations in the plant community composition data, where the maximum height was less than the median height; 4) LDMC values at or outside the bounded interval 0−1; 5) SLA values below <0 or >200 mm²/mg⁻¹; 6) Leaf area, fresh mass, and dry mass values of ≤0; 7) Observations in the plant functional trait data, where dry mass exceeded fresh mass; 8) Image-derived colour indices we checked against their theoretical ranges: −1-1 for ExG and 0-1 for BITM. Based on these criteria, we inspected all flagged observations and set the traits to missing when we confirmed that there were errors or if we were not able to locate and correct the source of the error.

When inspecting the data, we noticed that a simple statistical outlier detection based on univariate methods is poorly suited to our ecological trait data collected along wide environmental gradients. This is because the extreme values often reflect genuine biological variations, and also, because the distributions are typically right-skewed and highly species-specific. We therefore based the main automated detection procedure on the within-species covariance structure of functionally related trait pairs. This approach rests on the principle that most of the traits are not independent: leaf wet and dry weight share a near-constant ratio within a species, and leaf area scales predictably with both wet and dry weight. Consequently, their derivatives (.ie., LDMC and SLA) are also typically highly correlated. Severe single-observation deviations from these expected relationships within a species are therefore more indicative of measurement error than of true biological variation.

Next, we focused on four trait pairs: log(d_weight) against log(w_weight), log(leaf_area) against log(d_weight), log(leaf_area) against log(w_weight), and log(SLA) against LDMC. For these pairs, we fitted a robust linear regression per species using Huber M-estimation as implemented in the *rlm* function of the *MASS* package (Venables and Ripley 2002). Here, we preferred robust regression over ordinary least squares because the latter is sensitive to the very outliers we seek to detect: a single erroneous observation can pull the fitted line toward itself, reducing its own residual and evading detection. The Huber estimator down-weights observations with large residuals during fitting, so the resulting line reflects the central tendency of the majority of observations and outlying points to receive appropriately large residuals. Because all continuous trait distributions were right-skewed and spanning several orders of magnitude, prior to fitting, we applied log-transformation for all pairs except LDMC against SLA, where LDMC was left on its original scale as a bounded near-symmetric variable. Here, we included only species with ≥10 complete observations for a given pair. We studentised residuals from each robust fit by dividing them with their median absolute deviation (MAD), yielding a dimensionless measure of how many MADs each observation lies from the species-specific regression trend. Then, we retained this MAD-studentised residual as a continuous diagnostic variable for each trait pair.

Since the three weight and leaf area pairs share raw trait measurements (i.e., leaf area, wet weight, dry weight), we could use the pattern of high residuals across the pairs to identify which specific raw measurement is most likely in error. This is because each of the three raw trait measurements is absent from exactly one of the three pairs: leaf area does not appear in the dry-against-fresh-weight pair, dry weight does not appear in the area-against-fresh-weight pair, and fresh weight does not appear in the area-against-dry-weight pair. Consequently, if a measurement was erroneous, the one pair that did not involve it would show a low residual while the two pairs that did involve it would show high residuals. We used this logic to assign each observation a suspect variable: observations that had a high residual in the leaf area pairs but a low residual in the weight pair indicated a suspicious leaf area; observations that had a high residual in the dry-weight pairs combined with a low residual in the wet-weight-area pair indicated a suspicious dry weight; and the complementary pattern indicated a suspicious fresh weight. If observations did not fit any of these three clean patterns, we classified them as ambiguous and we inspected them manually. We used the SLA–LDMC pair as a confirmatory signal rather than a diagnostic one, since either or both of these derived traits are affected if any of the raw trait measurements is erroneous. We subjected to diagnosis observations with a MAD residual >5 in any of the three raw trait measurement pairs, and subsequently, we set the traits to missing according to the identified suspect variable and its downstream consequences for derived traits: a suspicious leaf area propagated to missing SLA, BITM, and ExG; a suspicious fresh weight propagated to missing LDMC; and a suspicious dry weight propagated to missing SLA and LDMC. Additionally, we observations to missing if they had a MAD residual >8 in the SLA–LDMC pair that had not been resolved by the pattern-based diagnosis, and we did this across all derived trait measurements and raw trait measurements, as deviations of this magnitude indicated a measurement inconsistency too severe to attribute to a single source. In total, we determined 14 leaf samples containing clearly erroneous trait values and we set their particular traits to missing.

## Usage notes

### Data use and best practice notes

We provide this dataset under a CC-BY license. We recommend users of the dataset to cite this data article when referencing and using these data. We encourage contact from users for guidance, advice, and collaboration. We appreciate users contacting us before visiting the study sites due to our long-term monitoring programs.

### Strengths

A major strength of this dataset is the high level of operator consistency. This means that all field and laboratory work was conducted by the same two researchers. Therefore, we were able to limit subjectivity in this dataset, reducing errors that may arise from observer bias and maximising comparability across study designs and study areas.

A second major strength is the nested plot structure that we used in many of the study designs, enabling multi-level analyses. However, this structure also requires careful consideration, because the trait measurements represent different levels of observation. This means that we documented the plant community composition and plant height at the plot-level, whereas, we measured leaf traits and reproductive effort at the site-level.

Finally, a third key strength is the high spatial accuracy of the recorded study site coordinates. We provide coordinates up to centimetre-scale positioning accuracy, ensuring that the exact same study sites can be reliably relocated in the future, and that the spatial context of the observations incorporated into analyses. High positioning accuracy also enabled us to do true resurveys of the exact same sites, plots, and plant communities in the Kilpisjärvi Saana-Jehkas gradient design, Kilpisjärvi Community experiment design, and Kilpisjärvi Seasonal monitoring design.

### Considerations

Temporal context is one the most important factors to consider when using this dataset. We collected the data over more than a decade. This means that different parts of the dataset originate from different years, both within and among study areas and study designs. During our observation and sampling period 2016–2025, the climatic conditions in northern Europe were extreme at times, including winter warming and snow-on-ice events (Aalto et al. 2026). Moreover, 2024 was likely the warmest growing season in northern Europe in the past 2000 years (Rantanen et al. 2025).

Seasonal dynamics should also be considered. We collected the data during peak growing seasons, except in the Kilpisjärvi Seasonal monitoring design, in which we documented seasonal patterns in leaf traits, see Niittynen et al. (2026). However, growing season dynamics can vary substantially in northern ecosystems. For instance, the onset of the growing season may shift between years and vary across local environmental gradients (Rantanen et al. 2026).

Land use practices and particularly grazing pressure contribute to variation among study areas and study sites. For example, the study sites in Kilpisjärvi study area are located within three distinct reindeer pastures. Thus, the grazing pressure by the semi-domesticated reindeer (*Rangifer tarandus tarandus*) and its spatiotemporal dynamics vary greatly even within the study area. Reindeer grazing affects the plant community composition and likely also plant heights (Olofsson et al. 2009, Maliniemi et al. 2018, Happonen et al. 2019).

The dataset captures a wide range of environmental variation within study areas, because we selected the study sites within each study area using stratified random sampling. However, this dataset should not be considered an unbiased representation of entire landscapes, and this should be considered especially when calculating species-level average trait values. For example, atypical habitats are overrepresented relative to their true frequency, such as springs in the Kilpisjärvi Spring design and geofeatures in the Kilpisjärvi Geodiversity gradient design.

Furthermore, we measured vegetative height excluding inflorescences. However, plant height measurements are dependent on the presence of inflorescence in many species. In our study areas, species such as *Solidago virgaurea* often occurred only with the leaf rosettes close to the ground, but their height measurements were considerably taller, if they produced flowers at the top of a tall shoot with many leaves.

Regarding leaf traits, we also measured the leaf traits of horsetails (*Equisetum* spp.). However, using these data calls for consideration because these species do not really have leaves. Therefore, we used the smaller branches as equivalents of leaves for some species, such as *E. sylvaticum*. Whereas, for the non-brancing species (*E. scirpoides, E. variegatum, E. hyemale*, and *E. fluviatile*), we calculated the leaf traits using the whole green shoot.

Finally, we emphasise that the plant community composition data and the plant height data represents the entire plant community of each site, whereas the leaf trait data does not. This is because we collected leaf samples of the entire community at each site, except for species with a low coverage ≤ 1%. More importantly, we did not sample any protected species, in the countries in which they were protected. Therefore, the leaf trait data do not represent the entire plant community.

### Limitations and uncertainties

Regarding taxonomy, we consider the main source of uncertainty to be species misidentification. We mitigated this risk through extensive prior experience in species identification of these northern ecosystems (Niittynen et al. 2018, Kemppinen et al. 2021a). In addition, all leaf sample identifications were verified by both of us during laboratory work. Nevertheless, we acknowledge that hybridisation is common in certain groups, such as *Salix* spp. and *Carex* spp., and such cases we have explicitly marked as hybrids in the dataset.

Regarding the plant community composition data, we consider the main limitation to be the visual estimation that we used for estimating the species coverages. We minimised observer bias because all estimates were made by the same two researchers, and we cross-calibrated our estimations. However, visual estimation is inherently more subjective than, for instance, image-based analysis or the point-intercept method. As a result, direct comparisons of species cover with future resurveys should be conducted with particular care.

Regarding plant height, we consider the main source of uncertainty to be the median height which we visually estimated. This means that we first visually inspected the representative median height of the given species in the plot, and then based on this inspection, we measured an individual representative of median height. We chose this approach to reduce workload, but a more robust approach would have involved measuring a larger number of individuals and then calculating the median based on their measured heights. Consequently, additional measurements would be required to obtain more objective estimates of median height.

Finally, regarding leaf trait data, we consider the main limitation to be that we pooled the leaf samples at site-level. We chose this approach to reduce workload, but a more comprehensive approach would have involved conducting the leaf trait measurements at plot-level and individual-level. Our approach limits the ability to quantify variation within-plot and within-population. Consequently, more detailed sampling at finer scales and sampling more local replicates would be required to reduce potential noise arising from inter-individual and intra-individual variation which affects the trait averages at plot-level (Maitner et al. 2023).

## Author contributions

Conceptualisation (equal), Data curation (equal), Formal analysis (PN), Funding acquisition (equal), Investigation (equal), Methodology (equal), Project administration (equal), Resources (equal), Software (PN), Supervision (equal), Validation (PN), Visualization (equal), Writing - original draft (equal), Writing - review & editing (equal)

## Acknowledgements

We thank the personnel at the following research stations for their support during our fieldwork and laboratory work: Kilpisjärvi Biological Station and Värriö Subarctic Research Station, Oulanka Research Station, and Kevo Subarctic Research Institute. We thank the University of Helsinki, University of Oulu, and University of Turku, our work would not be possible without these invaluable research stations across northern Finland. We thank Berit Tønsberg Gaski and Kari Anne Bråthen for their support during fieldwork at Máttavárri and giving us access to the Climate-ecological Observatory for Arctic Tundra cabin. We thank Tuuli Rissanen for sharing species lists of the Rásttigáisa study sites with us. We thank Johanna Lehtinen and Miska Luoto for sharing species lists of the Pallas study sites with us. We thank Ian Brown from Stockholm University for helping to establish the Vindelfjällen study design. We thank the 4th Plant Functional Trait Course held in Svalbard 2018, particularly Vigdis Vandvik, Aud H. Halbritter, Brian Maitner, and Brian J. Enquist who taught us how and why to sample plant functional traits.

## Permits

Permission to carry out fieldwork was granted by Metsähallitus.

## Supporting information

**Figure S1.**
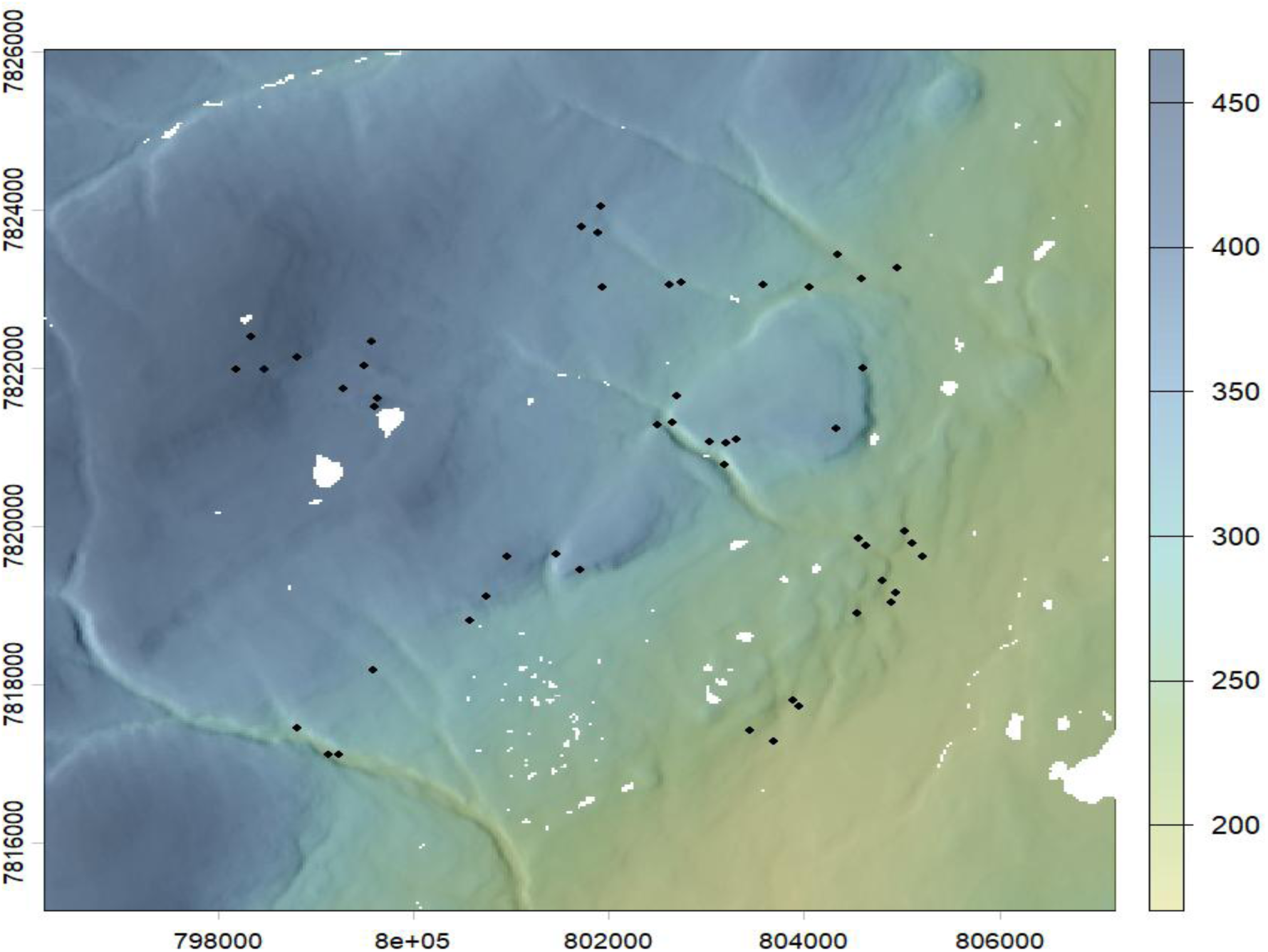
Map of the Máttavárri study area. Colours represent elevation (m a.s.l.) overlaid with hillshade. Black dots represent the study site locations. Water bodies in white. The coordinate system is UTM34N WGS84.

**Figure S2.**
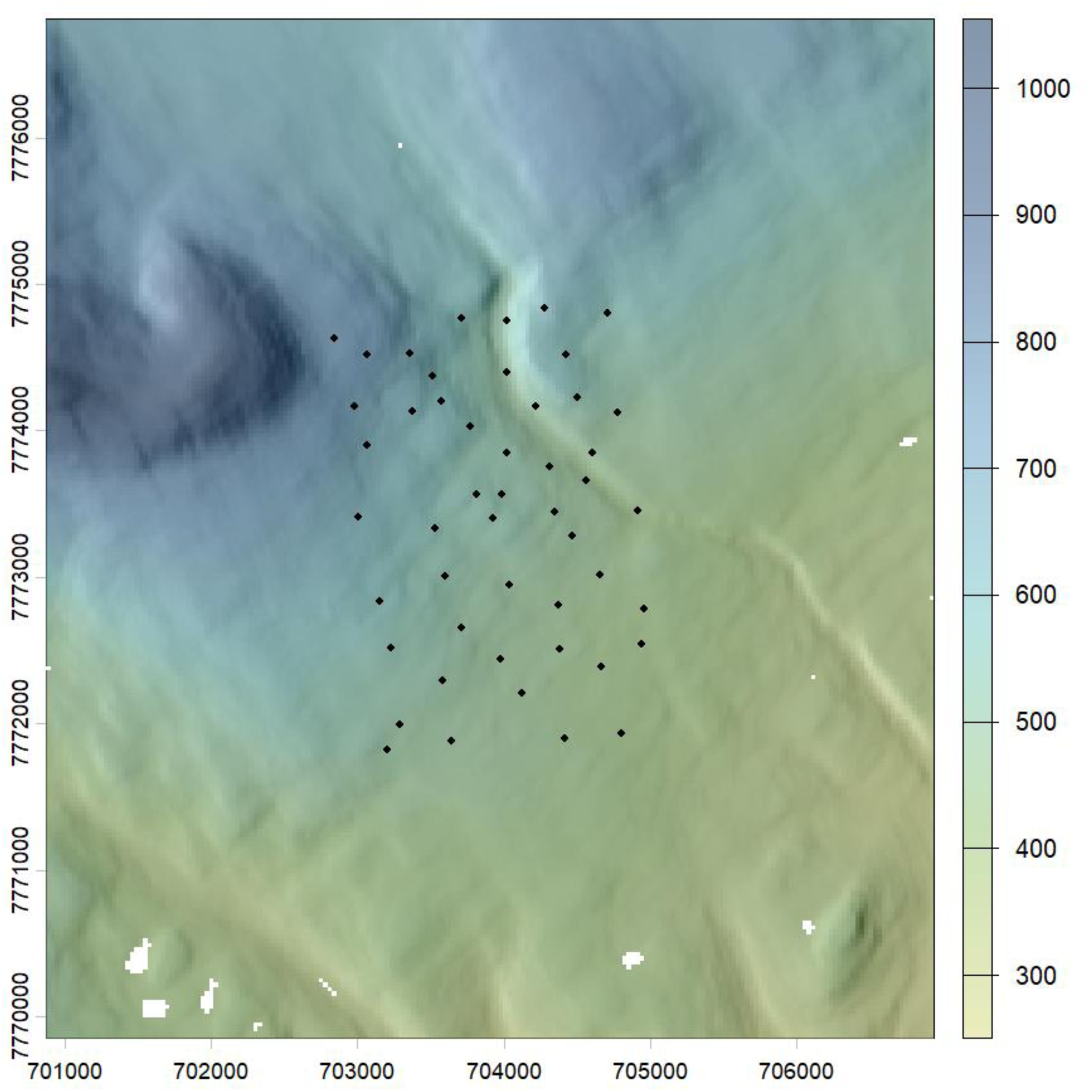
Map of the Rásttigáisá study area. Colours represent elevation (m a.s.l.) overlaid with hillshade. Black dots represent the study site locations. Water bodies in white. The coordinate system is UTM34N WGS84.

**Figure S3.**
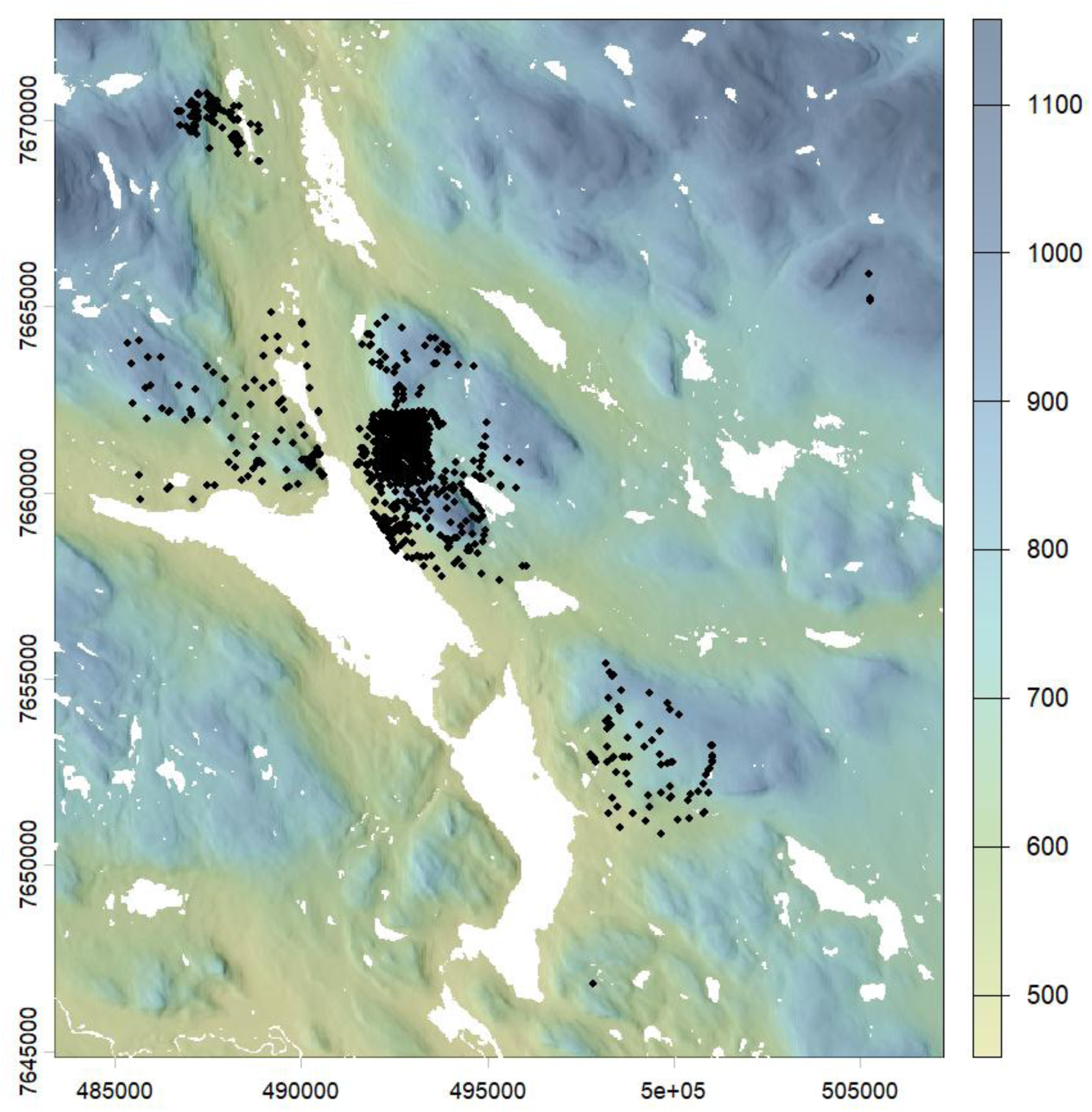
Map of the Kilpisjärvi study area. Colours represent elevation (m a.s.l.) overlaid with hillshade. Black dots represent the study site locations. Water bodies in white. The coordinate system is UTM34N WGS84.

**Figure S4.**
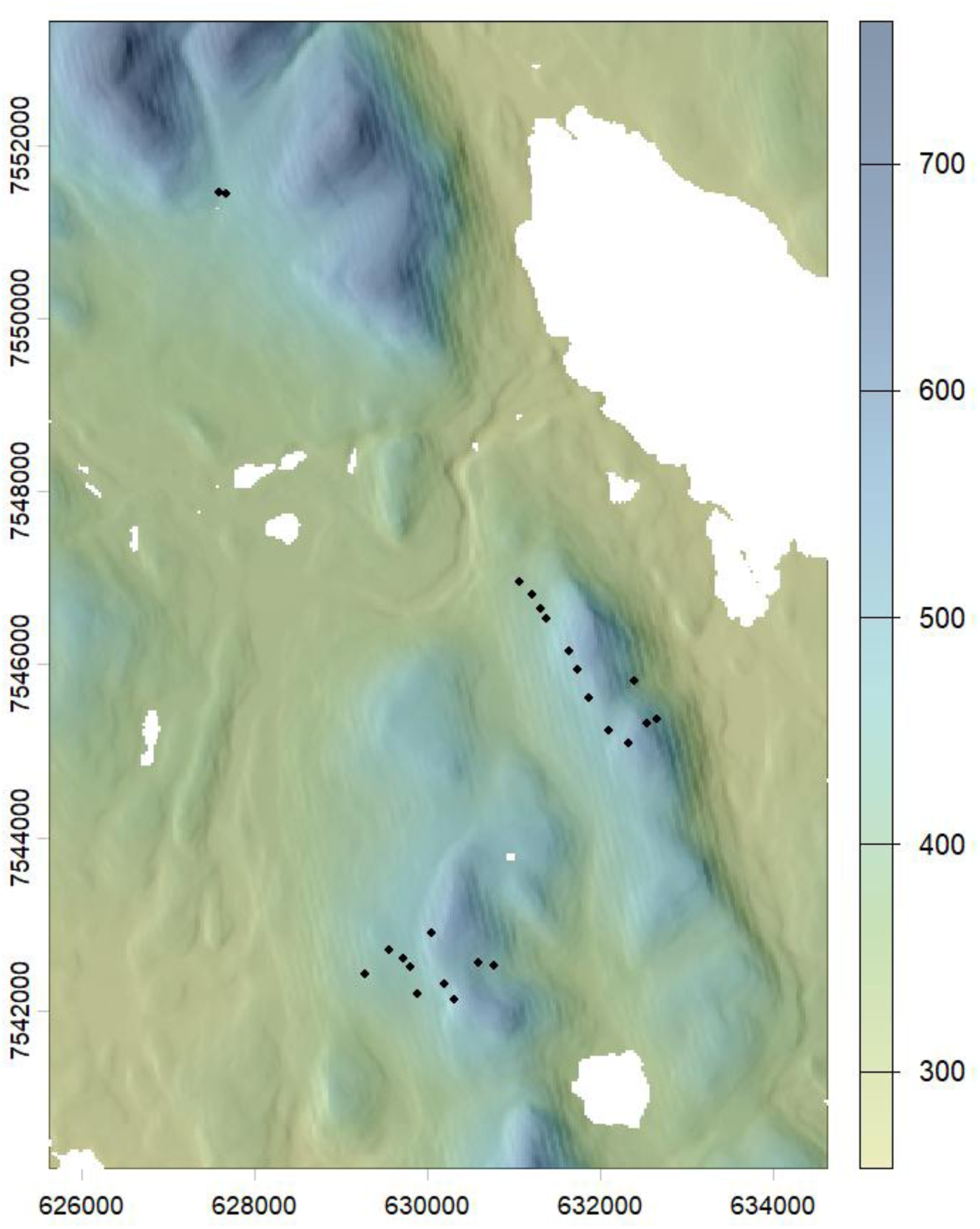
Map of the Pallas study area. Colours represent elevation (m a.s.l.) overlaid with hillshade. Black dots represent the study site locations. Water bodies in white. The coordinate system is UTM34N WGS84.

**Figure S5.**
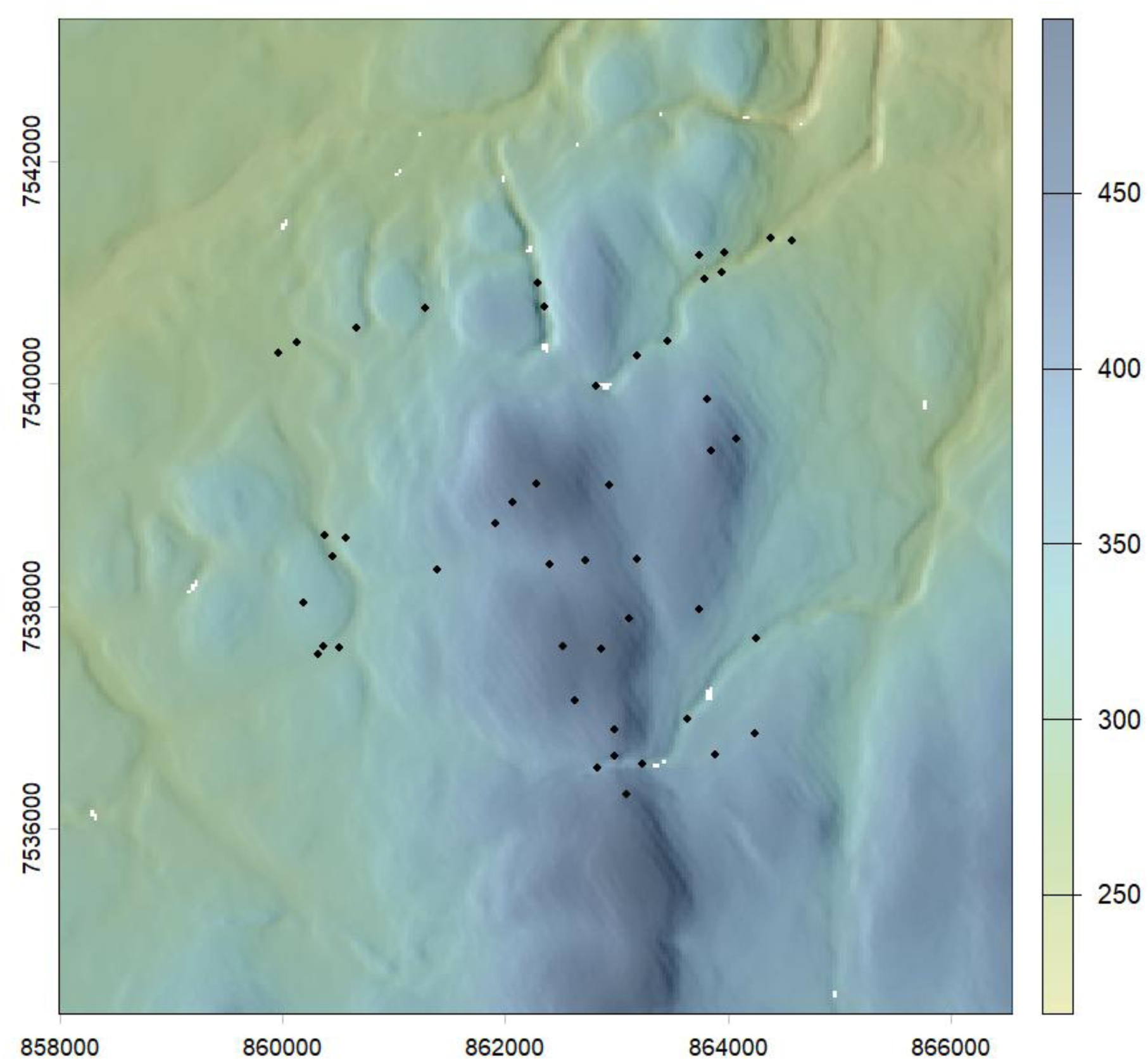
Map of the Värriö study area. Colours represent elevation (m a.s.l.) overlaid with hillshade. Black dots represent the study site locations. Water bodies in white. The coordinate system is UTM34N WGS84.

**Figure S6.**
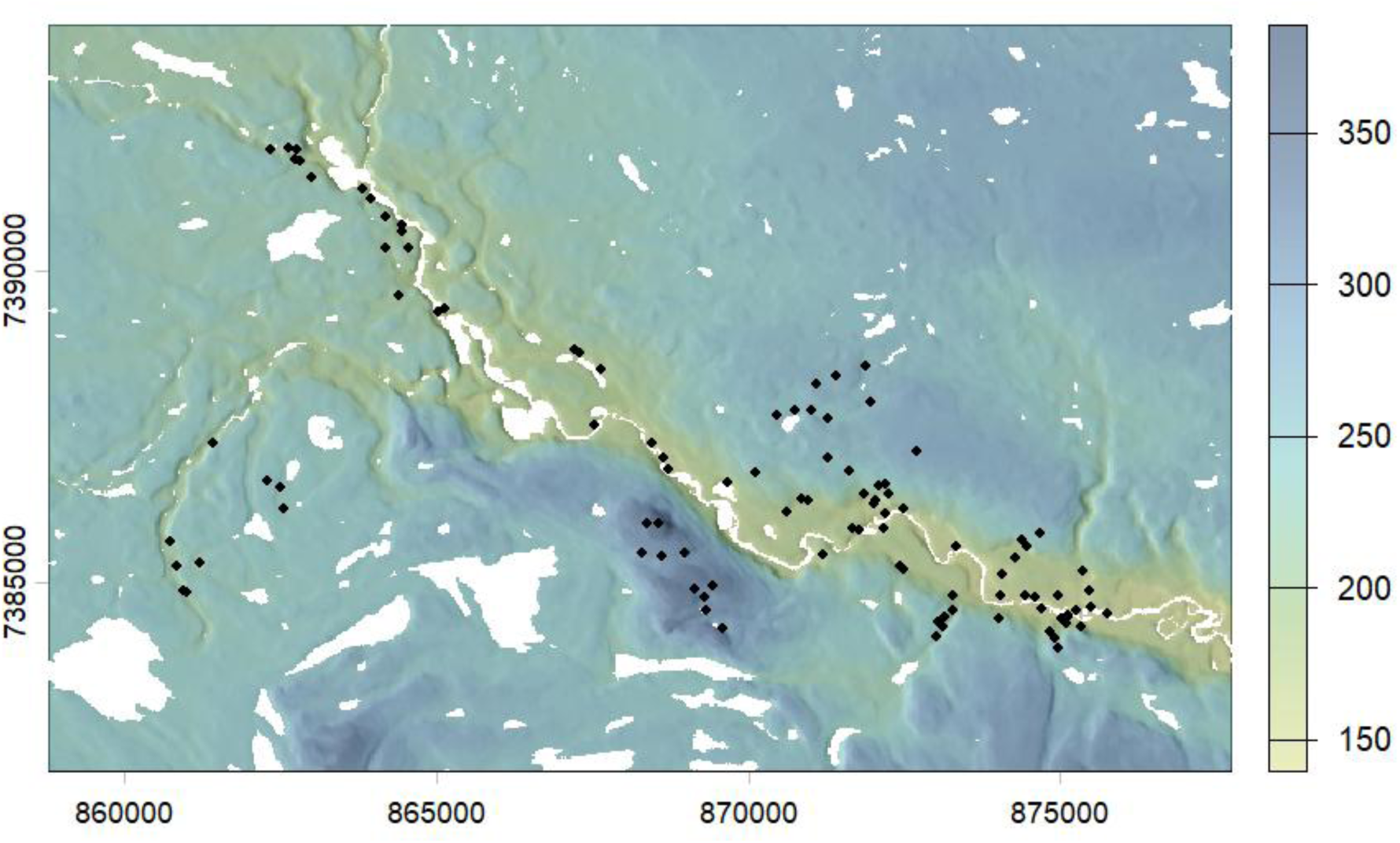
Map of the Oulanka study area. Colours represent elevation (m a.s.l.) overlaid with hillshade. Black dots represent the study site locations. Water bodies in white. The coordinate system is UTM34N WGS84.

**Figure S7.**
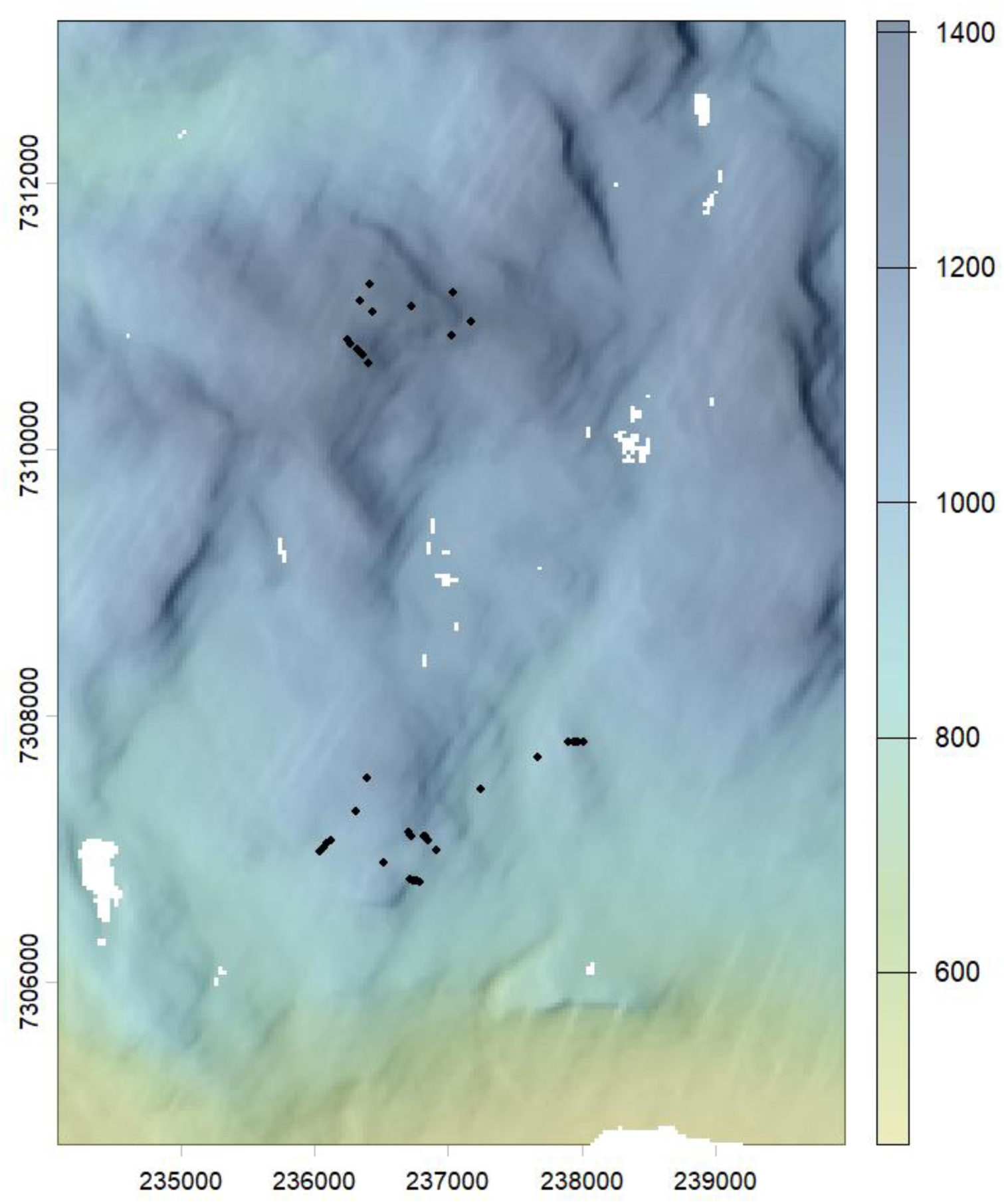
Map of the Vindelfjällen study area. Colours represent elevation (m a.s.l.) overlaid with hillshade. Black dots represent the study site locations. Water bodies in white. The coordinate system is UTM34N WGS84.

