## Supplementary material for "FennoTraits: Dataset of plant functional traits and community composition in northern European flora": FennoTraits_combine_data.R

```
#######################################################################
# This R script facilitates the start in working 
# with the *FennoTraits* data
#######################################################################

# Load tidyverse packages
library(tidyverse)

# read in the community and height data
comm <- read_csv("data/FennoTraits_community_heights.csv")
# read in the leaf and reproductive trait data
leaf <- read_csv("data/FennoTraits_leaf_traits.csv")
# read in the taxonomic lookup table
tax <- read_csv("data/FennoTraits_species.csv")

# Take a glance on the datasets
comm %>% summary()
leaf %>% summary()
tax %>% summary()

# Combine the trait datasets into a wide-format data table
# Note! Heights are at plot level but other traits at site level. 
# Thus those other traits are replicated to all plots within each site.
all <- full_join(comm %>% 
                   select(site:date, taxon:max_height),
                 leaf %>% 
                   select(site:date, taxon, reproduction, leaf_area:ExG)) %>% 
  left_join(.,
            tax %>% select(taxon, rank)) %>% 
  relocate(rank, .after = taxon)


# Drop taxa at higher taxonomic level or is a hybrid
all <- all %>% 
  filter(!rank %in% c("hybrid","genus","family"))

# Filter only 1-m plot data with full communities surveyed and no experimental manipulation
all <- all %>% 
  filter(plot_type == "b", # b indicates the 1-m plots
         full_community == TRUE,
         treatment == "none")
  
# Transform community composition data to site x species matrix
cm <- all %>% 
  mutate(taxon = gsub(" ", "_", taxon)) %>% # get rid of spaces in the to-be column names
  pivot_wider(id_cols = c(site, design, area, date), names_from = taxon, values_from = cvr) %>% 
  mutate(across(-c(site, design, area, date), ~replace_na(.x, 0))) # replace NA's with zeros
str(cm)

# Get a long-format trait table
all_long <- all %>% 
  pivot_longer(cols = median_height:ExG, names_to = "trait", values_to = "value", values_drop_na = TRUE)

# Plot abundance distribution of Empetrum nigrum
cm %>% 
  ggplot(aes(x = Empetrum_nigrum)) +
  geom_histogram(bins = 20)+
  theme_bw()

# Plot all trait distributions of Viola biflora
all_long %>% 
  filter(taxon == "Viola biflora") %>% 
  ggplot(aes(x = value)) +
  geom_histogram(bins = 20)+
  theme_bw() +
  facet_wrap(vars(trait), scales = "free")
```
